# Thermal motion of DNA in an MspA pore

**DOI:** 10.1101/021766

**Authors:** Bo Lu, Stephen Fleming, Tamas Szalay, Jene Golovchenko

**Affiliations:** Department of Physics, Harvard University, Cambridge MA 02138; School of Engineering and Applied Sciences, Harvard University, Cambridge Massachusetts, 02138, USA

## Abstract

We report on an experiment and calculations that determine the thermal motion of a voltage-clamped ssDNA-NeutrAvidin complex in an MspA nanopore. The electric force and diffusion constant of DNA inside an MspA pore have been determined in order to evaluate DNA’s thermal position fluctuations. We show that an out-of-equilibrium state returns to equilibrium so quickly that experiments usually measure a weighted average over the equilibrium position distribution. Averaging over the equilibrium position distribution is consistent with results of state-of-the-art nanopore sequencing experiments. It is shown that a reduction in thermal averaging can be achieved by increasing the electrophoretic force used in nanopore sequencing devices.

## INTRODUCTION

The protein nanopore MspA, with its relatively sharp and narrow pore constriction, has provided a very promising platform for sequencing DNA(1-6). To measure a sequence-specific signal, single-stranded DNA (ssDNA) would ideally remain in a clamped state (see Fig.1A, top) created by two opposing forces: (1) the electrophoretic force driving DNA through the nanopore, and (2) an opposing force provided by a molecular stop, such as an enzyme(7), a region of double-stranded DNA (dsDNA)(2), or any bound protein(6). The several nucleotides near the narrowest constriction of MspA control the passage of ionic current through the pore, providing a current measurement corresponding to the nucleotides in the constriction region. One might then expect the length of the narrowest constriction of MspA to completely determine how many nucleotides contribute to the measured current signal. However, previous studies show that the actual length of DNA sampled during a current measurement is longer that the length of the pore’s constriction(1, 2, 6). This raises an interesting question: Do thermal fluctuations of DNA inside the pore affect the nucleotide resolution of the device, and can these fluctuations be minimized?

For a short piece of DNA trapped in a pore using a molecular stop, the same thermal fluctuations that affect the positional sensitivity for sequencing can also cause the molecule to stochastically escape from the nanopore entirely (Fig. 1B, bottom). The distribution of escape times can be modeled as a process of one-dimensional, diffusive escape from an energy trap using the first-passage formalism. Using this model, we extract values for the electrophoretic driving force and the diffusion constant from measurements of the escape time distributions at a range of clamping voltages. These parameters have previously been reported for the α-hemolysin nanopore using various experimental measurements and models(8-13), including approaches similar to ours(14). Even though the nucleotide discrimination regions of α-hemolysin have been mapped out(15), the 12-base-long beta-barrel constriction of α-hemolysin still makes it difficult to distinguish the mixing due to thermal motion from a purely geometric averaging effect. Thus in previous work, using the obtained electrophoretic driving force and diffusion constant for the evaluation of DNA thermal motion has not attracted significant attention.

**Figure 1.**
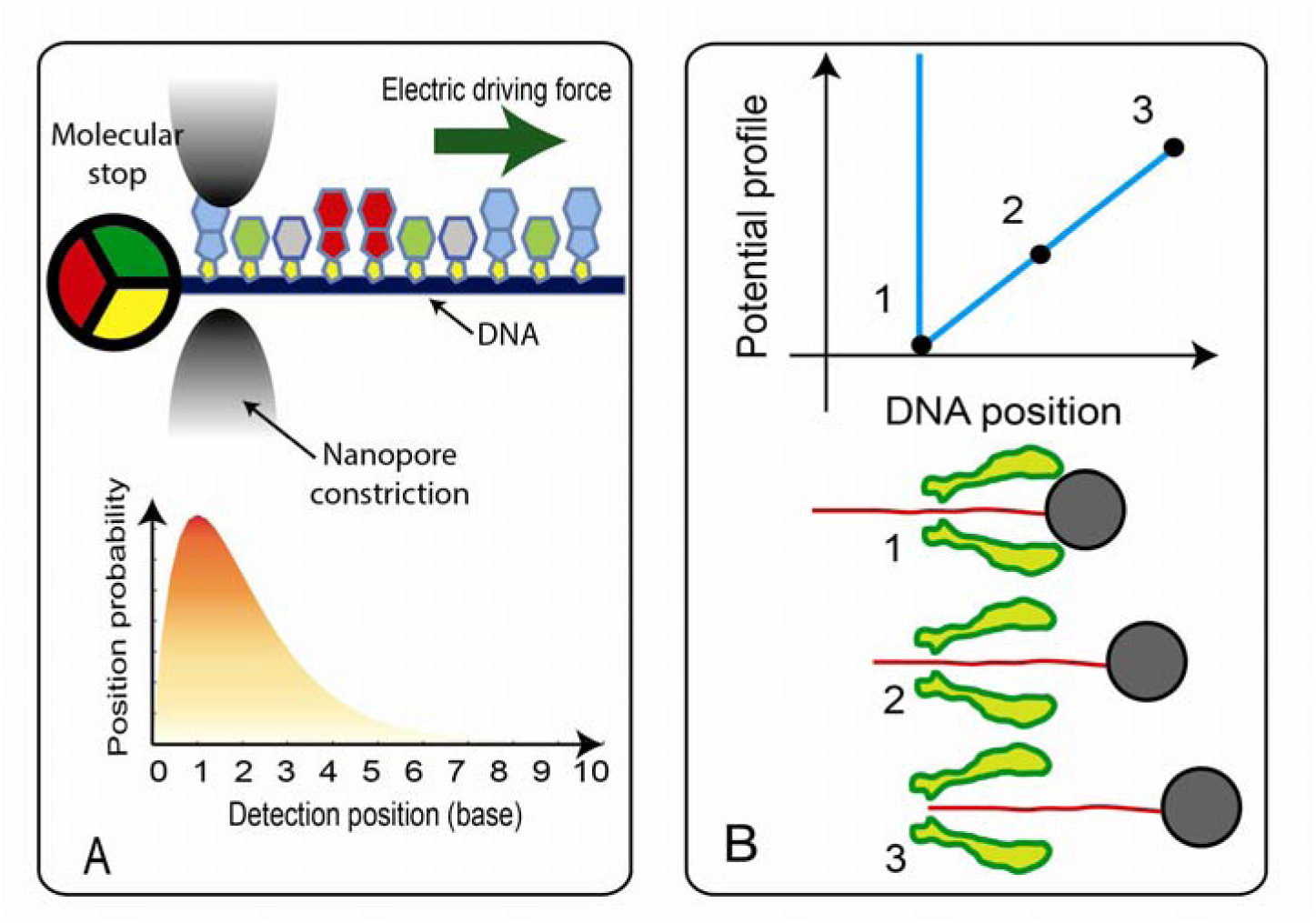
(A) In this schematic diagram, ssDNA is trapped inside a nanopore by a molecular stop, which opposes an electrophoretic driving force. For nanopore sequencing strategies, thermal motion’s impact on nucleotide resolution can be described by a position probability function, which depends on the driving force and diffusion constant of DNA. (B) The energy landscape for a trapped ssDNA in an MspA pore. The gray sphere is an attached NeutrAvidin molecule that prevents the ssDNA from translocating through the pore. The escape rate and time for such trapped DNA depends on the driving force and diffusion constant of DNA.

In this work, we measure escape times of a ssDNA-NeutrAvidin complex from MspA at low clamping voltages in order to calculate the thermal fluctuations of DNA under typical clamped sequencing conditions, and we show that these fluctuations can produce a weighted average over the bases near the constriction (Fig. 1A). Such fluctuations have a direct impact on the observation that while approximately 2 nucleotides fit into MspA’s narrowest constriction(1, 2), nearly 4-nucleotide resolution has been demonstrated in previous MspA work(4, 6). Briefly, we model DNA motion as a one-dimensional diffusion process in a simplified energy trap landscape with a reflecting boundary and a sloping energy barrier created by the electrophoretic driving force (see Fig. 1B). The electrophoretic driving force on DNA and the diffusion constant of DNA in an MspA pore is then extracted by fitting a first-passage model to the experimental escape times. These parameters are used to evaluate the equilibrium position probability distribution as well as the time-dependent evolution toward equilibrium, which corresponds to sequencing conditions. We find that the evolution of the position probability density from a non-equilibrium state to equilibrium is much faster than 1 µs. Within our instrument bandwidth, the position fluctuations contribute to the measured current signal as a weighted average of position-dependent current over the equilibrium position distribution. This “thermal averaging” decreases the resolution of the MspA pore. We determine that for typical sequencing conditions, 99% of the position probability distribution is spread over a 2∼3-nucleotide region, and support this with previous MspA pore sequencing data showing the footprint of a single-nucleotide mutation in a poly-dA region, and we discuss how nucleotide resolution can be improved.

## EXPERIMENTAL METHOD

### ssDNA-NeutrAvidin complex escape experiment

The detailed procedure of the ssDNA-NeutrAvidin escape experiment is as follows. DNA was first captured in an MspA pore by applying 160mV capture voltage. This voltage is chosen based on the requirement that the ssDNA-NeutrAvidin complex must not escape backward out of the pore. Our measurements show that poly-dA, with the 5’ end leading, blocks 75.5% of total pore conductance, which is slightly smaller than previous reports of 72.1%,(6) but the value is reasonable considering this data is obtained using a different mutant of MspA. 20ms after a complex is trapped in the pore, the driving voltage is switched to a constant, low value. While the driving voltage is low (35mV-70mV), DNA captured by the nanopore (Fig. 2A) can more easily escape (as shown in Fig. 2), and cannot translocate through the pore due to the large size of the NeutrAvidin molecule. The escape time of the DNA is defined as the duration from the moment the voltage switches to the low state to the moment the pore current returns to the open, unblocked state (as shown in Fig. 2C).

**Figure 2.**
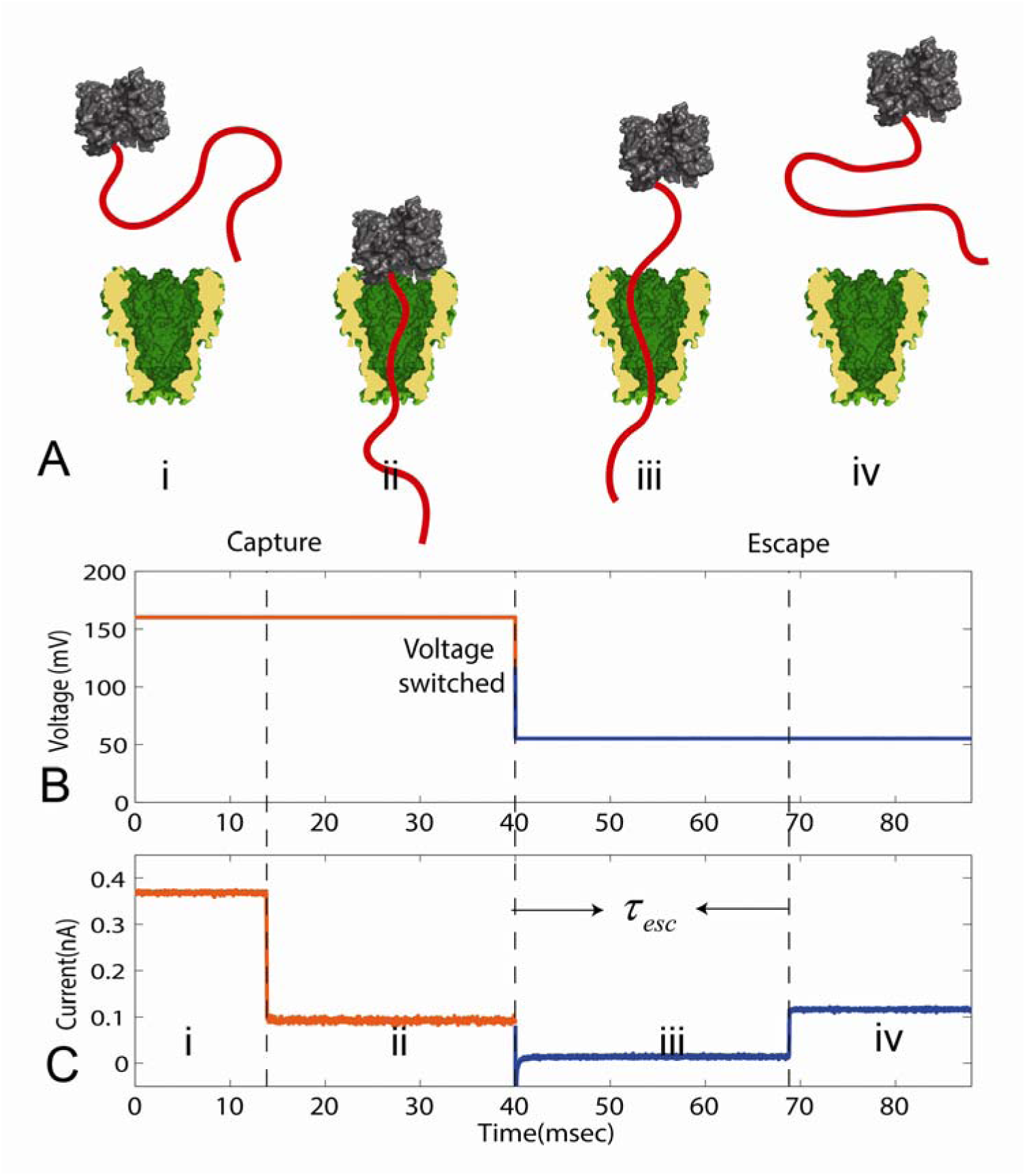
(A) Schematic figure of a ssDNA-NeutrAvidin complex being captured (i, ii) and escaping (iii, iv) from an MspA pore. (B), (C) The dynamic voltage and current trace demonstrate the capture and escape process. The definition of escape time has been labeled in (C).

## EXPERIMENTAL MATERIAL

The MspA protein nanopore was provided by Oxford Nanopore Technologies, Inc., and is the G75S/G77S/L88N/D90N/D91N/D93N/D118R/Q126R/D134R/E139K mutant of wild-type MspA. The methods for forming a single MspA nanopore in a lipid bilayer membrane are described in several papers(1, 16, 17). Briefly, a lipid membrane composed of 1,2-diphytanoyl-*sn*-glycero-3-phosphocholine [DPhPC] (Avanti Polar Lipids, Inc.) is formed on a 30 micron diameter Teflon aperture, separating two reservoirs filled with a buffer solution containing 1M KCl, 10mM HEPES at pH 8.0. Hexadecane was used as the hydrophobic lipid solvent which creates an annulus connecting the lipid bilayer to the Teflon support. A voltage bias was applied across the lipid membrane using Ag/AgCl electrodes. During experiments, the measured current and voltage were low-pass filtered at 10kHz and recorded at a sampling frequency of 250kHz using an Axopatch 200B (Molecular Devices, Sunnydale, CA). A single pore insertion was characterized by approximately 2.3nS conductance at 23?, ambient room temperature.

DNA was purchased from IDT with HPLC purification. The DNA used in these experiments was a single strand of poly-dA, 27 nucleotides in length, with a 3’ biotin. ssDNA was mixed with NeutrAvidin (Thermo Scientific) in a 1:3 (ssDNA:NeutrAvidin) molar ratio by slowly adding ssDNA to a concentrated NeutrAvidin solution while stirring, in order to ensure that most NeutrAvidin proteins had only one ssDNA bound. Approximately 10pM NeutrAvidin-DNA complex was loaded into the reservoir where the ground electrode is connected.

The voltage control setup is based on an Arduino microcontroller with a digital-to-analog output converter. The Arduino is programmed to react to certain conductance conditions by adjusting the command voltage appropriately, which it sets using the voltage input of the Axopatch 200B.

## RESULT AND DISCUSSION

### Determination of electrophoretic driving force and diffusion constant

By repeating this capture and escape process mentioned in Fig. 2, a distribution of escape times can be obtained, as shown in Fig. 3A-C. Considering the 10kHz low-pass filter setting, events shorter than 100μs have not been included in the distribution plot and further data analysis. The escape time distribution takes the form of an exponential, at least for short times that encompass about 80% of the total number of escape events. For longer time scales, the escape time distribution follows a power law(14). According to the first-passage formalism, DNA escape from an energy trap should follow an exponential escape time distribution. Furthermore, the average escape time’s dependence on the clamping voltage can be understood in the context of a one-dimensional first-passage approach. In this analysis, the non-exponential region has been excluded from the average escape time distribution. The detailed data analysis procedure and further explanation of the non-exponential tail of the escape time distribution are contained in the appendix.

**Figure 3.**
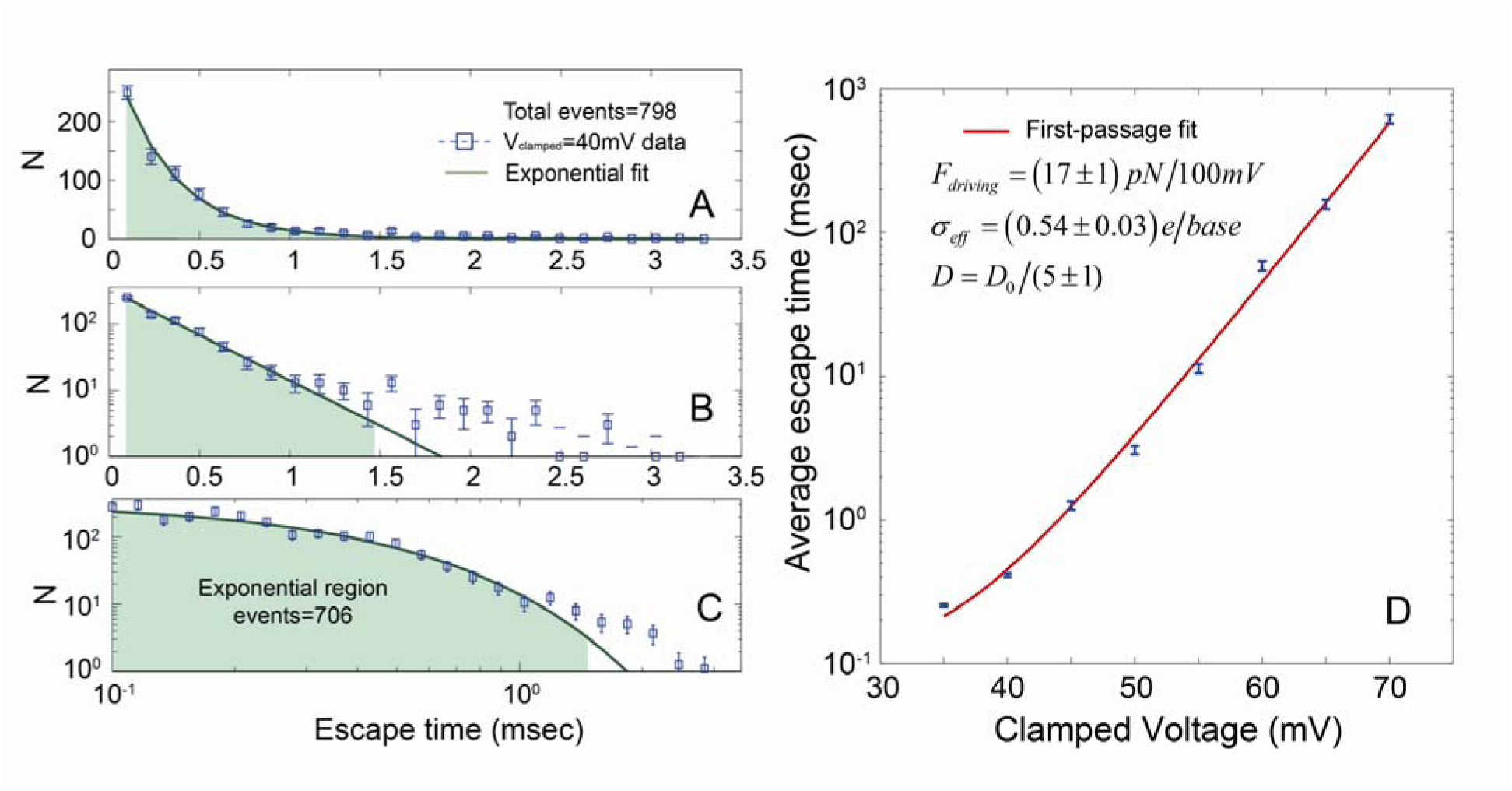
(A) Escape time distribution for ssDNA at 40mV clamping voltage. The same escape time distribution is plotted on log scale axis in (B) and (C) to show the exponential decay profile at short times. (D) Plot of the average escape time as a function of clamping voltage. The red curve is the first-passage calculation fit. *D*_*0*_ is defined as the diffusion constant for a 1.5nm diameter sphere in free space.

As shown in Fig. 3D, the average escape time can be plotted as a function of the clamping voltage from 35mV to 70mV. We fit the curves with a first-passage model in order to extract the electrophoretic driving force and diffusion constant, as described in our previous work(18). The point where MspA’s narrowest constriction meets the ssDNA captured inside the pore is defined as a “walker,” diffusing along a domain of *x* ∈[0, *L*_*domain*_]. *x =* 0 is the position of MspA’s constriction along the ssDNA when the NeutrAvidin is pulled against the entrance to the pore’s vestibule, and *x* = *L*_*domain*_ is the position of MspA’s constriction along the ssDNA when the end of the ssDNA tail escapes from the constriction altogether. By considering the length of DNA(19) and the total size of the MspA pore(6, 20), *L*_*domain*_ is set to be the length of 14 nucleotides, with length per base *a*_0_ =0.5nm. According to the first-passage approach, we define the probability of passage times as *f*_*esc*_ (*x*_0_, *t*) *dt*, which represents the probability that the DNA escapes from the pore back into solution within a time between *t* and *t + dt,* given an initial starting position *x*_*0*_. This probability function obeys the adjoint Smoluchowski equation:

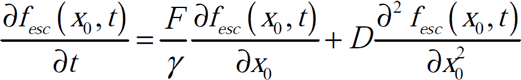

An absorbing boundary is set at the end of the ssDNA tail, *f*_*esc*_(*x*_0_, *t*)|_*x*_0_=*L*_*domain*__ = *δ* (*t*), representing the end of an escape process. The repulsive force caused by the NeutrAvidin’s attempt to enter the vestibule of the nanopore can be represented in a simplified manner as a reflecting boundary at *x =* 0: 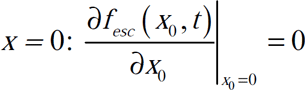. This boundary condition is based on the assumption that the electrophoretic driving force used in the experiment is not strong enough to detach NeutrAvidin from the 3’ biotin linker on the ssDNA, an assumption which previous experiments have proven valid(6). The initial condition is *f*_*esc*_ (*x*_0_,0)= 0, (*x*_0_ < *L*_*domain*_). Here *D* and *γ* are the diffusion constant and drag coefficient, which are related through the fluctuation-dissipation theorem. The force applied to DNA in this domain can be written as *F* = *F*_*V*_ + *F*_*entropy*_. This force includes the electrophoretic driving force, *F*_*V*_, which is commonly defined as an effective charge density multiplied by the potential drop across the nanopore(21, 22), *F*_*V*_ = *σ* _*eff*_ *V*. The total force also includes an entropic force that is attributable to the conformational freedom of the ssDNA, *F*_*entropy*_. *F*_*entropy*_ is negligible on most of the domain, except when only the very tail end of the ssDNA is inside the constriction region of the pore. The entropic force would then be *F*_*entropy*_ = *k*_*B*_*T* / *l*_*Kuhn*_, for *L*_*domain*_ - *l*_*Kuhn*_ ≤ *x* ≤ *L*_*domain*_, where *l*_*Kuhn*_ is the Kuhn length of ssDNA, approximately 1.5nm(16, 19). Other than that, the entropic effect can be neglected since the entropy does not vary much over the rest of the domain. Hence, the only free parameters in the adjoint Smoluchowski equation are the effective charge, *σ*_*eff*_, and the diffusion constant, *D*.

The escape time distribution is given by 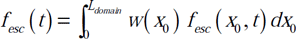. Here, *w* (*x*_0_) is the spread of initial positions of the first-passage walker. Initial position obeys equilibrium exponential distribution *w*(*x*_0_) = exp (*F*_*C*_*x*_0_/*k*_*B*_*T*) *F*_*C*_*/k*_*B*_*T*, where *F*_*C*_ = *σ* _*eff*_ *V*_*capture*_ is the driving force under capture voltage 160mV. DNA’s average escape time from the nanopore is given by 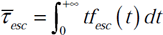. To be consistent with the time limit imposed by the 10kHz filter, the integral limit is rectified to 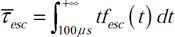. Note that because diffusive motion dominates the escape process, the initial position of the walker actually has little effect on the escape time distribution, as long as the initial position is far away from the escape boundary at *x* = *L*_*domain*_. This insensitivity to initial conditions shows that the capture voltage hardly affects the escape time distribution. The first-passage model is optimized by using a least-squares fit with two free parameters: *σ*_*eff*_and *D*. The best fits for 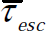 are shown as the red curves in Fig. 3D. The parameter values extracted in this manner are *σ* _*eff*_ = 0.54 ± 0.03*e* / *base*, and *D* = *D*_0_ / (5 ± 1), where *D*_0_= 2.94 ×10^-10^ *m* ^2^*/s* is the diffusion constant of a 1.5nm diameter bead free in solution, 1.5nm being the approximate Kuhn length of ssDNA. We account for the uncertainty in the length per base by using the value 0.5 ±0.02nm(23), which leads to a 4% relative uncertainty in the effective charge density.

### Evolution of the DNA position distribution

Using our measurement of the electrophoretic driving force and the diffusion constant, we calculate the position probability function to evaluate DNA’s position uncertainty under typical experimental sequencing conditions.

The domain of the position probability function is the same as for the first-passage approach: *x* ∈[0, *L*_*domain*_]. The position probability function, *P* (*x*, *x*_0_, *t*), is defined for a walker with a starting position *x*_0_at *t* = 0 as the probability of finding this walker at position *x* at time *t*. The position probability function obeys the Smoluchowski equation:

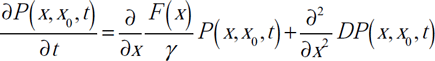

To examine the time-dependence of position fluctuations, we impose an initial condition that the ssDNA starts from position *x*_0_= 4*a*_0_, which can be expressed as *P*(*x*, 4*a*_0_,0) = *δ* (*x* − 4*a*_0_). The electrophoretic driving force pushes the ssDNA to approach the boundary at *x* = 0, where a simplified, reflecting boundary condition 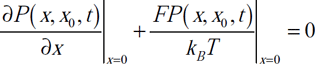 represents an infinitely high energy barrier created by the fact that the NeutrAvidin molecule is too large to enter the vestibule of the pore. If the electrophoretic driving force is large enough, the walker should have a low probability of approaching the boundary at *x* = *L*_*domain*_, signifying escape. This can be written as *P* (*x*, *x*_0_, *t*)_*x*_=*L*_*domian*_= 0.

The position probability function, plotted in Fig. 4A, shows that an initial displaced delta function will develop into the equilibrium exponential distribution in less than 10*n*sec. Such a fast relaxation time indicates that during the sampling interval we use to make a measurement of the current (100∼250kHz sampling rate and 10kHz lowpass filter), the DNA has enough time to explore all the possible positions under the equilibrium distribution. Or in other words, our instrument actually measures a weighted average of position-dependent current over the entire equilibrium position probability distribution.

**Figure 4.**
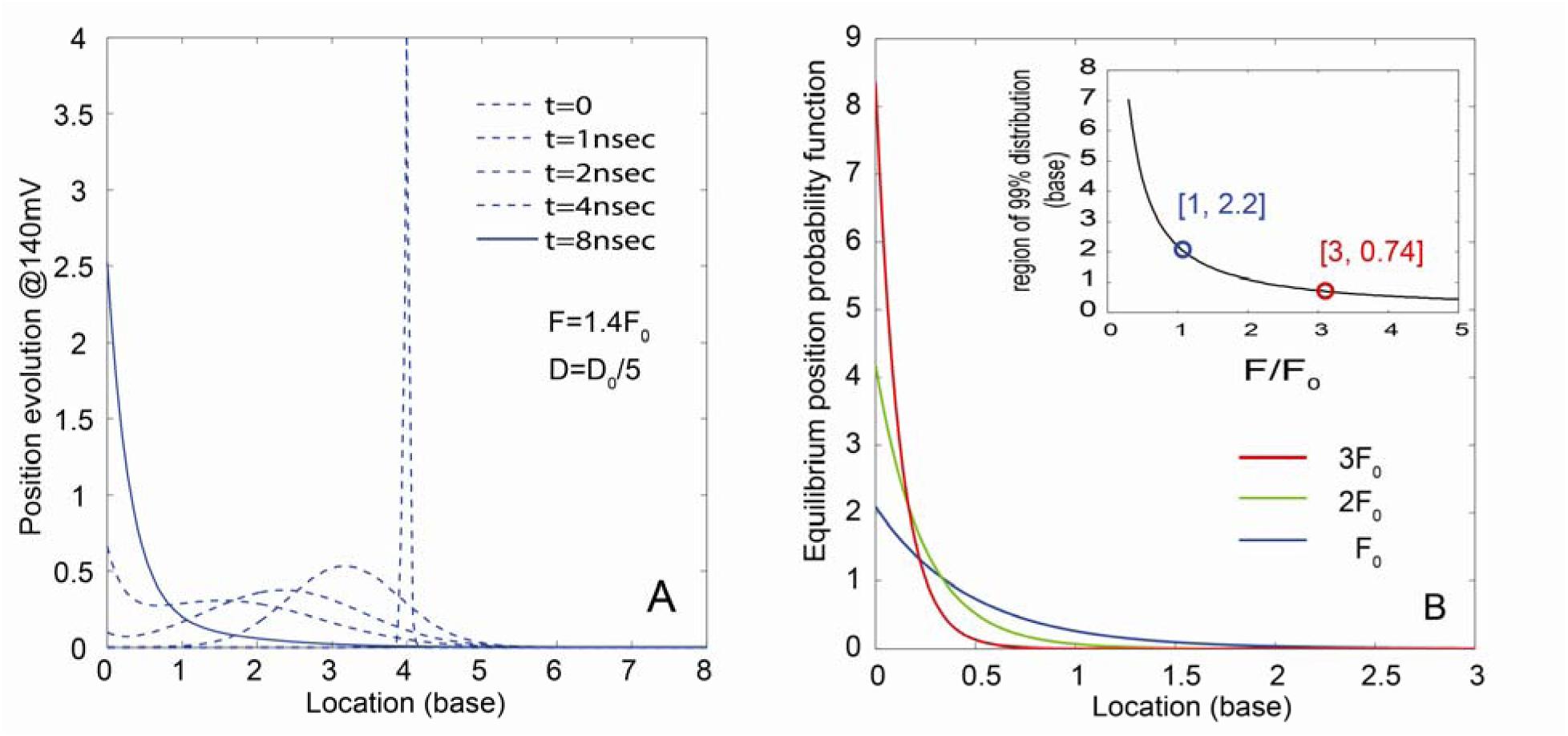
(A) The normalized DNA position probability function evolves from a delta function to an equilibrium exponential function within 10nsec at 140mV driving voltage. *F*_*0*_ is 17pN, defined as the driving force for 0.54e/base charge density ssDNA at 100mV. *D*_*0*_ is defined as the diffusion constant for a 1.5nm diameter sphere in free space. (B) The equilibrium position probability function under different electrophoretic driving forces. The inset shows the region encompassed by 99% of the position distribution function, and its dependence on the driving force.

The relaxation time mentioned in Fig. 4A can be understood intuitively using a back-of-the-envelope calculation. A position fluctuation caused spontaneously by thermal motion needs some time to be restored to the equilibrium distribution, and this corresponds to the elaxation time. Assuming the thermal fluctuation perturbs the position of the pore’s constriction relative to the ssDNA from its original *x* = 0position, where the equilibrium probability *p* = exp(−*Fx /k*_*B*_*T*) *F / k*_*B*_*T* is *p* = *F kBT*, to a position *x* = *k*_*B*_*T F*, where the equilibrium probability is *p* = exp(−1) *F k*_*B*_*T*, then the time the driving force needs to restore the displacement is greater than the time scale given by 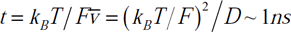, which is equivalent to the relaxation time for the given position fluctuation.

DNA position fluctuations can cause uncertainty when mapping nanopore sequencing data onto an actual DNA sequence. Typically, nanopore sequencing experiments measure a series of stable current “levels” which correspond to the bases in the constriction region, and the durations of these levels vary. According to the previous calculation, mapping these measured current levels to sequence information only presents a problem in the case where the duration of a current level is shorter than the aforementioned relaxation time, since in that case, the current measurement collected by the instrument will correspond to a local region covering only part of the equilibrium position distribution. Position fluctuations that occur on time scales shorter than the relaxation time are irregular and unpredictable. However, the sampling rate of our instrument, and instruments typically used for sequencing experiments, is about 10us. The typical duration of a current level in a nanopore sequencing experiment is about 5-10ms or even longer(7). The levels last much longer than the relaxation time for position fluctuations, which means that the current averaged over the entire level duration is actually the average of position-dependent current over thermal position fluctuations.

The relaxation time for DNA thermal fluctuations depends on the magnitude of the electrophoretic driving force and the diffusion constant of DNA in the nanopore. Note that the diffusion constant measured in the α-hemolysin pore is about 300 times(12) or 2000 times(13) smaller than that measured in this work. One might expect that the diffusion constant could be larger in MspA than in α-hemolysin due to differences in pore geometry, with the long, narrow constriction of α-hemolysin leading to a smaller diffusion constant. Additionally, the large difference can also be attributed to the fact that these measurements of the diffusion constant in the α-hemolysin pore were for hairpin DNA instead of NeutrAvidin-tethered ssDNA. Using our experimental setup and MspA, we measured that the diffusion constant of hairpin DNA is 250 times smaller than the one we report for the ssDNA-NeutrAvidin complex. We posit that the duplex of hairpin DNA has a more complicated interaction with the pore, being constrained inside the pore vestibule. Previous research has mentioned that positive charges in the MspA vestibule can enhance the capture capability of the pore(1), and we expect they might interact with the duplex of a hairpin as well. Additionally, the driving force for poly-dA in MspA may depend on the specific pore mutant. It is also natural to expect that the driving force or diffusion constant could differ for different homopolymer ssDNA, due not only to the identity of the different bases but also the differences in secondary structure.

So far, regardless of all the possible factors that could impact the diffusion constant and electrophoretic driving force, one conclusion is unchanged: The relaxation time for DNA thermal fluctuations in an MspA pore is much shorter than the time scale over which a sequence-specific current signal is measured. And a key point for future work is that DNA sequencing experiments could suppress thermal fluctuations in DNA’s position by using a larger driving force. This is true for all protein pores and solid-state pores. Fig. 4B shows that a larger driving force can make the position probability distribution narrower. We define the driving force of 0.54e/base with 100mV across the pore as the unit force *F*_*0*_, and quantify how the spread of the region encompassing 99% of the position probability distribution depends on driving force. This is shown in the inset of Fig. 4B. As the driving force becomes three times larger than *F*_*0*_, the position fluctuations will mainly be confined to a 1-nucleotide region.

We expect the thermal position fluctuations we have evaluated for clamped ssDNA in MspA to be highly relevant for nanopore sequencing applications. In previous MspA pore reports(2, 4-6), the number of nucleotides contributing to a single current level measured in an MspA pore was estimated to be about 3 or 4 at 180mV bias. The "nucleotide resolution" of the device can be characterized by the current change caused by a single heteromeric substitution in homopolymer ssDNA. As shown by Laszlo *et al.(4)*, reproduced here in Fig. 5, the presence of a single G substitution in a poly-dA strand has a large effect on ionic current when the G substitution is in a specific 2-base region, since GAAA and AGAA deviate from AAAA by 23%. Here the sequence denotes which bases are in the constriction of MspA, leftmost being the first through the pore. As the G substitution moves to AAGA and AAAG, the current value deviates from AAAA by 3.7%. Beyond this 4-base region, the substitution’s impact on the current is reported to be negligible. This data is in agreement with the limits on nucleotide resolution imposed by the position probability function we calculated in Fig. 4B, where under 180mV driving voltage, 99% of the distribution curve is spread over a 1.3-base region, not including the first base located at *x* = 0. We expect that as we update the reflecting boundary to a repulsive force *F*_*barrier*_(*x*) the position probability distribution should spread slightly beyond *x* = 0. This repulsive force would have a finite magnitude over a certain region due to effects like DNA stretching(23).

**Figure 5.**
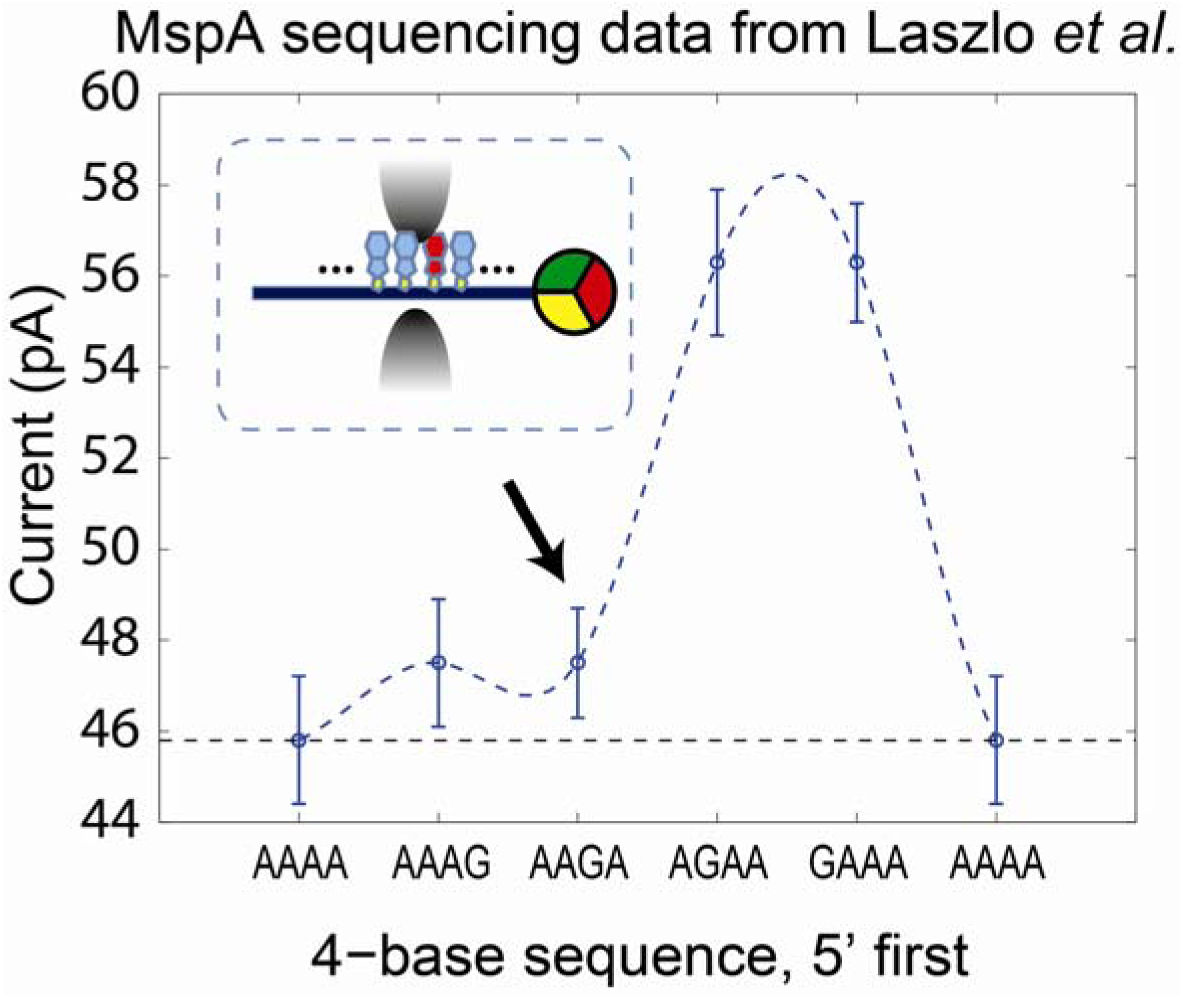
Data taken from Laszlo et al., a previous report about MspA sequencing data. Plot shows the current signal corresponding to a 4-mer sequence with a G substitution at different locations. The error bar on each point is the standard deviation of the mean current level for the specific 4-mer sequence measured at different positions along phi X 174 DNA. The inset shows the orientation of AAGA inside the MspA pore.

State-of-the-art nanopore sequencing strategies rely on the measurement of ionic current through a voltage-biased nanopore as a DNA molecule is ratcheted, in single base steps, through the pore by discrete enzymatic action.(7) Position fluctuations of the enzyme itself during the phase of the ratcheting process when the enzyme is stationary on the DNA could in turn contribute to position fluctuations of DNA. These thermal motions ultimately provide a limit on the nucleotide resolution of nanopore sequencing devices. Other limits result from the chemostochastic timing of the ratcheting enzyme and the finite number bases influencing the ionic current even in the absence of thermal fluctuations. In addition, the resolution can be further impacted by nucleotide-pore interactions, which remain uncharacterized.

## CONCLUSION

Using the first-passage formalism, the effective charge and diffusion constant of ssDNA in an MspA nanopore have been extracted from a ssDNA-NeutrAvidin escape experiment. The position probability function for ssDNA in MspA based on these two parameters shows that the stochastic dynamics of DNA manifest as high frequency noise, beyond the sampling rate of our current amplifier. For the time scales typically used to characterize a current level during sequencing experiments, the thermal motion averages the signal over an extended domain of a few nucleotides. Such thermal averaging inevitably contributes to the nucleotide resolution of the device; however, comparisons with nucleotide resolution in previous MspA work show that thermal motion is likely only one of several important factors that determine the overall nucleotide resolution. The magnitude of the electrophoretic driving force required to optimize the nucleotide resolution of an MspA nanopore device has been identified. In the future, other sources of position fluctuations such as DNA-pore interactions and enzyme ratcheting action will require further investigation in order to better interpret nanopore sequencing data and develop devices with improved sequencing accuracy.

## APPENDIX

Analysis of the exponential decay region in the escape time distribution

The exponential region can be extracted as follows. First, an R-squared fit to the line f(X) = k_l_X+ k_2_ is performed on a histogram of escape events with a log scale y-axis. The R-squared value was used to indicate the goodness of the fit. We started by using only the short time region (40% of total events) to do the exponential fit. Then we kept increasing the amount of data, and we optimized the bin size used for the distribution in order to obtain the best R-square value. The parameters corresponding to the best R-square value determine the optimal value of k_l_and the region that can be best fit by the exponential curve. Using these fits, the time domain which spans 99% area under the exponential curve exp (k_l_X) is plotted as the exponential region shown as green in Fig. 3A,B and C. All the escape events too long to be included in this exponential region have been excluded in the average escape time calculation. The nonexponential region is beyond the scope of what can be predicted by the first-passage model. The power law distribution on long time scales may be caused by irregular interactions between DNA and nanopore, which are still in need of further investigation.

## ACKNOWLEDGMENTS

We thank Professor Daniel Branton for his advice on the experiment, and Eric Brandin for his assistance with sample preparation and the experiment. This research was supported by National Institutes of Health award HG003703. Stephen Fleming was supported by the National Science Foundation Graduate Research Fellowship under Grant No. DGE1144152.

## References

1. Butler, T. Z., M. Pavlenok, I. M. Derrington, M. Niederweis, and J. H. Gundlach. 2008. Singlemolecule DNA detection with an engineered MspA protein nanopore. Proceedings of the National Academy of Sciences 105:20647–20652.

2. Derrington, I. M., T. Z. Butler, M. D. Collins, E. Manrao, M. Pavlenok, M. Niederweis, and J. H. Gundlach. 2010. Nanopore DNA sequencing with MspA. Proceedings of the National Academy of Sciences 107:16060–16065.

3. Laszlo, A. H., I. M. Derrington, H. Brinkerhoff, K. W. Langford, I. C. Nova, J. M. Samson, J. J. Bartlett, M. Pavlenok, and J. H. Gundlach. 2013. Detection and mapping of 5-methylcytosine and 5-hydroxymethylcytosine with nanopore MspA. Proceedings of the National Academy of Sciences 110:18904–18909.

4. Laszlo, A. H., I. M. Derrington, B. C. Ross, H. Brinkerhoff, A. Adey, I. C. Nova, J. M. Craig, K. W. Langford, J. M. Samson, R. Daza, and J. H. Gundlach. 2014. Decoding long nanopore sequencing reads of natural DNA. Nature biotechnology.

5. Manrao, E. A., I. M. Derrington, A. H. Laszlo, K. W. Langford, M. K. Hopper, N. Gillgren, M. Pavlenok, M. Niederweis, and J. H. Gundlach. 2012. Reading DNA at single-nucleotide resolution with a mutant MspA nanopore and phi29 DNA polymerase. Nature biotechnology 30:349–353.

6. Manrao, E. A., I. M. Derrington, M. Pavlenok, M. Niederweis, and J. H. Gundlach. 2011. Nucleotide discrimination with DNA immobilized in the MspA nanopore. PLoS One 6:e25723.

7. Cherf, G. M., K. R. Lieberman, H. Rashid, C. E. Lam, K. Karplus, and M. Akeson. 2012. Automated forward and reverse ratcheting of DNA in a nanopore at 5-A precision. Nature biotechnology 30:344–348.

8. Sauer-Budge, A. F., J. A. Nyamwanda, D. K. Lubensky, and D. Branton. 2003. Unzipping kinetics of double-stranded DNA in a nanopore. Physical Review Letters 90:238101.

9. Mathé, J., H. Visram, V. Viasnoff, Y. Rabin, and A. Meller. 2004. Nanopore unzipping of individual DNA hairpin molecules. Biophysical Journal 87:3205–3212.

10. Lathrop, D. K., E. N. Ervin, G. A. Barrall, M. G. Keehan, R. Kawano, M. A. Krupka, H. S. White, and A. H. Hibbs. 2010. Monitoring the escape of DNA from a nanopore using an alternating current signal. Journal of the American Chemical Society 132:1878–1885.

11. Lakatos, G., T. Chou, B. Bergersen, and G. N. Patey. 2005. First passage times of driven DNA hairpin unzipping. Physical biology 2:166.

12. Wanunu, M., B. Chakrabarti, J. Mathé, D. R. Nelson, and A. Meller. 2008. Orientation-dependent interactions of DNA with an α-hemolysin channel. Physical Review E 77:031904.

13. Mathé, J., A. Aksimentiev, D. R. Nelson, K. Schulten, and A. Meller. 2005. Orientation discrimination of single-stranded DNA inside the α-hemolysin membrane channel. Proceedings of the National Academy of Sciences of the United States of America 102:12377–12382.

14. Wiggin, M., C. Tropini, V. Tabard-Cossa, N. N. Jetha, and A. Marziali. 2008. Nonexponential kinetics of DNA escape from α-hemolysin nanopores. Biophysical journal 95:5317–5323.

15. Stoddart, D., A. J. Heron, E. Mikhailova, G. Maglia, and H. Bayley. 2009. Single-nucleotide discrimination in immobilized DNA oligonucleotides with a biological nanopore. Proceedings of the National Academy of Sciences 106:7702–7707.

16. Smith, S. B., Y. Cui, and C. Bustamante. 1996. Overstretching B-DNA: the elastic response of individual double-stranded and single-stranded DNA molecules. Science 271:795–799.

17. Akeson, M., D. Branton, J. J. Kasianowicz, E. Brandin, and D. W. Deamer. 1999. Microsecond time-scale discrimination among polycytidylic acid, polyadenylic acid, and polyuridylic acid as homopolymers or as segments within single RNA molecules. Biophysical journal 77:3227–3233.

18. Hoogerheide, D. P., F. Albertorio, and J. A. Golovchenko. 2013. Escape of DNA from a weakly biased thin nanopore: experimental evidence for a universal diffusive behavior. Physical review letters 111:248301.

19. Chi, Q., G. Wang, and J. Jiang. 2013. The persistence length and length per base of singlestranded DNA obtained from fluorescence correlation spectroscopy measurements using mean field theory. Physica A: Statistical Mechanics and its Applications 392:1072–1079.

20. Perera, A. S., H. Wang, T. B. Shrestha, D. L. Troyer, and S. H. Bossmann. 2013. Nanoscopic surfactant behavior of the porin MspA in aqueous media. Beilstein journal of nanotechnology 4:278–284.

21. Keyser, U. F., B. N. Koeleman, S. Van Dorp, D. Krapf, R. M. Smeets, S. G. Lemay, N. H. Dekker, and C. Dekker. 2006. Direct force measurements on DNA in a solid-state nanopore. Nature Physics 2:473–477.

22. Lu, B., D. P. Hoogerheide, Q. Zhao, and D. Yu. 2012. Effective driving force applied on DNA inside a solid-state nanopore. Physical Review E 86:011921.

23. Stoddart, D., L. Franceschini, A. Heron, H. Bayley, and G. Maglia. 2015. DNA stretching and optimization of nucleobase recognition in enzymatic nanopore sequencing. Nanotechnology 26:084002.

